# Gambogic acid and gambogenic acid induce a thiol-dependent heat shock response and disrupt the interaction between HSP90 and HSF1 or HSF2

**DOI:** 10.1101/2021.04.26.441566

**Authors:** Linda Pesonen, Sally Svartsjö, Viktor Bäck, Aurélie de Thonel, Valérie Mezger, Délara Sabéran-Djoneidi, Pia Roos-Mattjus

## Abstract

Cancer cells rely on heat shock proteins (HSPs) for growth and survival. Especially HSP90 has multiple client proteins and plays a critical role in malignant transformation, and therefore different types of HSP90 inhibitors are being developed. The bioactive natural compound gambogic acid (GB) is a prenylated xanthone with antitumor activity and it has been proposed to function as an HSP90 inhibitor. However, there are contradicting reports whether GB induces a heat shock response (HSR), which is cytoprotective for cancer cells and therefore a potentially problematic feature for an anticancer drug. In this study, we show that GB and a structurally related compound, called gambogenic acid (GBA), induce a robust HSR, in a thiol-dependent manner. Using heat shock factor 1 (*HSF1*) or *HSF2* knockout cells, we show that the GB or GBA-induced HSR is HSF1-dependent. Intriguingly, using closed form ATP-bound HSP90-mutants that can be co-precipitated with HSF1, a known facilitator of cancer, we show that also endogenous HSF2 binds to the HSP90-HSF1 complex. GB and GBA treatment disrupt the interaction between HSP90 and HSF1 and HSF2. Our study implies that these compounds should be used cautiously if developed for cancer therapies, since GB and its derivative GBA are strong inducers of the HSR, in multiple cell types, by involving the dissociation of a HSP90-HSF1-HSF2 complex.

## Introduction

HSP90 is an essential ATP-dependent molecular chaperone that is one of the most abundant proteins in eukaryotic cells (Johnson 2012). HSP90 has a vast repertoire of client proteins consisting of kinases and phosphatases, nuclear hormone receptors, actin and tubulin, and the proteasome subunits, and its activity and client specificity relies on different co-chaperones (Csermely et al. 1998; Pearl 2016). There are two isoforms of cytosolic HSP90 in humans; HSP90α is considered the main isoform that is induced upon stress, whereas HSP90β is only slightly inducible and more abundant than HSP90α under physiological conditions (Csermely et al. 1998). HSP90β is considered important for normal cellular processes such as differentiation and cytoprotection and for maintenance of the cytoskeleton (Sreedhar et al. 2004).

The heat shock response (HSR) is a universal stress protective pathway that is induced in response to proteotoxic stress, e.g. exposure to heat, proteasome inhibitors, and infections (Richter et al. 2010). The HSR is characterized by a fast and massive increase in the expression of molecular chaperones, such as the heat shock proteins (HSPs), which refold damaged proteins and prevent protein aggregation. The transcription of *HSP* genes is mediated by heat shock factors (HSFs). Upon stress, HSFs oligomerize and accumulate into the nucleus, and bind to specific heat shock elements (HSEs). During heat stress, hundreds of genes are upregulated and thousands are downregulated (Mahat et al. 2016; Vihervaara et al. 2017). HSF1 is regarded as the master regulator of the HSR in mammals whereas HSF2 is involved in differentiation and development (Joutsen and Sistonen 2019). HSF2 has, however, been shown to form heterocomplexes with HSF1 and modulate the expression of HSR genes, suggesting also a role in the HSR (Östling et al. 2007; Sandqvist et al. 2009).

According to the chaperone titration model, cytoplasmic HSF1 monomers are kept inert when complexed with chaperones, such as HSP70 and HSP90 (Gomez-Pastor et al. 2018). Today, it is still unclear whether HSF2 can also form complexes with chaperones. Upon protein-damaging stress, chaperones are required for folding of denatured proteins and HSF1 is released from the chaperone complex, trimerized and activated. The mechanism by which HSF1 is inactivated is not completely clear. HSF1 has been shown to be inactivated by distinct post-translational modifications and by negative feedback regulation by HSPs, in particular HSP70 and HSP40 (Kmiecik et al. 2020; Masser et al. 2020). Interestingly, HSP90 was also recently shown to bind to HSF1 trimers and to favor the release of HSF1 from HSEs (Kijima et al. 2018). Clearly, the interaction between HSF1 and HSP90 is multifaceted.

Due to the notion that many HSP90 clients have crucial roles in rapidly growing cancer cells, inhibition of HSP90 suppresses many signaling pathways that are important for cancer progression. A number of HSP90 inhibitors have been extensively studied, but none of these compounds have been approved for clinical use by the FDA (Yuno et al. 2018). The first isolated HSP90 inhibitors were geldanamycin and radicicol (Neckers and Workman 2012). These compounds were shown to bind to the N-terminal ATP-binding pocket of HSP90 by mimicking the conformation of ATP. However, although inhibiting HSP90 activity, these proved too toxic, insoluble, and metabolically unstable for clinical use (Neckers and Workman 2012). Several compounds have been synthesized using geldanamycin and radicicol as templates, including 17-AAG (17-allylamino-17-demethoxygeldanamycin, tanespimycin), 17-DMAG, STA-9090 and AUY-922 (Shrestha et al. 2016). These N-terminal HSP90 inhibitors, also called classic inhibitors, are the most studied inhibitors. However, most of them induce a HSR that stimulates the synthesis of HSP90, which is counterproductive when considering treating cancer (Wang and McAlpine 2015; Yuno et al. 2018).

Many bioactive natural compounds found in plants have been shown to inhibit HSP90 (Dal Piaz et al. 2015). Celastrol (tripterine), a pentacyclic triterpenoid derived from the plant thunder god vine (*Tripterygium wilfordii*), is an anti-inflammatory agent that has long been used in Chinese medicine to treat autoimmune and inflammatory diseases (Chen et al. 2018; Salminen et al. 2010). Celastrol has been shown to inhibit angiogenesis, migration and invasion, and to suppress cancer progression (Salminen et al. 2010), by affecting multiple targets in cancer cells, including the IKK-NF-κB pathway, HSP90, and the proteasome (Chen et al. 2018). Gambogic acid (GB) is a natural product derived from the gamboge resin of the *Garcinia hanburyi* tree, which similarly to celastrol, has also been used in Chinese medicine (Banik et al. 2018). GB is a polyprenylated xanthone with antitumor, antimicrobial, and anti-inflammatory effects (Banik et al. 2018; Kashyap et al. 2016). GB also suppresses the progression of many cancers *in vitro* and *in vivo* (Tang et al. 2017; Wu et al. 2004; Xia et al. 2017) and in China, it has been evaluated in a clinical trial focused on targeting advanced malignant cancers (Chi et al. 2013). GB, as celastrol, affects different target proteins in the cell, such as HSP90, NF-κB, c-Myc, PI3K, p-AKT, MDM2, and the proteasome (Banik et al. 2018; Kashyap et al.2016). Gambogenic acid (GBA) is another active ingredient of the resin of *Garcinia hanburyi* (Asano et al. 1996). GBA resembles GB but has a geranyl and a hydroxyl group instead of the ether ring in GB (Asano et al. 1996). GBA, like GB, is toxic to many different cancer cell lines (Huang, T. et al. 2019; Liu et al. 2017; Shen et al. 2020; Zhou et al. 2013). Whether GBA induces a HSR has not been studied before.

In this study, we show that acute treatment with GB and GBA induce a robust HSR in multiple cell types irrespective of their developmental origins. Furthermore, we establish that GB and GBA induce the HSR in a thiol-dependent manner. In addition, we show that GB or GBA treatment disrupt the protein-protein interaction between HSP90 and HSF1 and HSF2. The potential of GB and GBA to activate the HSF pathway could be detrimental in cancer therapies and should be carefully considered if using GB or GBA in treatments or for further drug development.

## Materials and methods

### Generation of *HSF1* knockout U2OS cells with CRISPR-Cas9

The human osteosarcoma U2OS cells (HTB-96, ATCC) *HSF1* knockout cells (*HSF1* KO) were generated with CRISPR-Cas9 as previously described for *HSF2* knockout U2OS cells (*HSF2* KO) (Joutsen et al. 2020). *HSF1* KO cell clones were genotyped by DNA sequencing of PCR products spanning the targeted region of the *HSF1* gene. The selected U2OS clone presented one single base insertion on *HSF1* exon 3 (Table 1), the sequence analysis of six independent PCR products shows the same mutation, suggesting that all the alleles have the same mutation. Guide RNA sequence targeting *HSF1* exon 3: 5’-TGTTATGTGCAGATGGCTTC-3’. The following primers were used for PCR for validation: forward (hHSF1_Cr_ex3_F): 5’-GGTCCTTGTGGGTATGAACCT-3’ and reverse (hHSF1_Cr_ex3_R): 5’-CACACTGGTCACTTTCCTCTTG-3’.

**Table 1.**
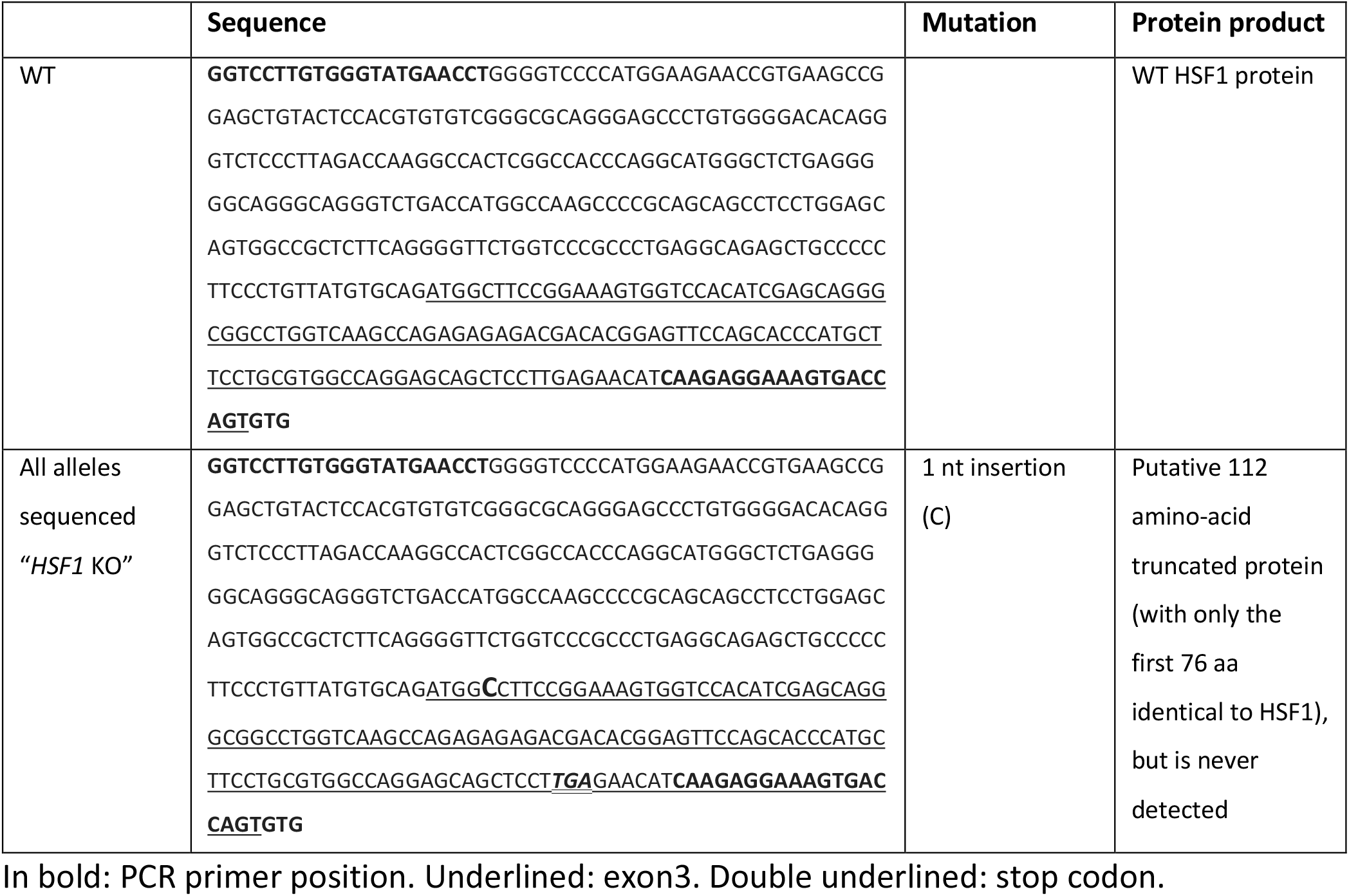
CRISPR-Cas9 induced mutation in HSF1 allele

### Cell culture and experimental treatments

HeLa (human cervical cancer, CCL-2, ATCC), HEK293 (human embryonic kidney cells, CRL-1573, ATCC), HDF (primary human dermal fibroblasts, PCS-201-010, ATCC), and U2OS (WT (HTB-96, ATCC), *HSF1* KO, and *HSF2* KO) cells were cultured in the same conditions: they were maintained at 37 °C in a humidified 5% CO_2_ atmosphere and cultured in Dulbecco’s modified Eagle’s medium (D6171, Sigma-Aldrich) supplemented with 10% fetal calf serum, 2 mM L-glutamine and 100 U/ml penicillin -100 µg/ml streptomycin mixture. RWPE-1 (human normal prostate epithelial cells, CRL-11609, ATCC) cells were cultured in Keratinocyte Serum Free Medium (17005042, Gibco) supplemented with 0.05 mg/ml bovine pituitary extract, 5 ng/ml epidermal growth factor, and 100 U/ml penicillin -100 µg/ml streptomycin mixture.

To induce a HSR and/or inhibit HSP90, cells were treated with either 17-AAG (17A, ant-agl-5, InvivoGen), celastrol (Cel, 3203, Tocris Bioscience), gambogic acid (GB, 3590, Tocris Bioscience) or gambogenic acid (GBA, BP2014, Chengdu Biopurify Phytochemicals Ltd.). The inhibitors were diluted in DMSO before treating the cells with concentrations indicated in the figures. Control cells were treated with DMSO only. Heat shock treatments were conducted on cell dishes wrapped in Parafilm and submerged in a water bath at 42 °C for indicated times. For the recovery phase, after treatments, the cells were placed in an incubator at 37 °C for 3 h after removing the parafilm. In order to investigate the thiol-reactiveness of GB and GBA, cells were, in addition to GB and GBA, also treated with dithiothreitol (DTT); GB + DTT and GBA + DTT, respectively. DTT was added in a 10-fold excess to GB and GBA, and left to react for 15 min at room temperature prior to cell treatment, with the indicated concentrations and the times, as described in the figure legends.

### Immunoblot analysis

Cells were lysed with either lysis buffer (25 mM HEPES pH 8.0, 100 mM NaCl, 5 mM EDTA pH 8.0, 0.5% Triton X-100) or Laemmli sample buffer (30% glycerol, 3% SDS, 187.5 mM Tris-HCl pH 6.8, 0.015% bromphenolblue, 3% β-mercaptoethanol). Cells lysed with lysis buffer were incubated in the buffer containing protease and phosphatase inhibitor cocktails (04693159001 and 04906845001, Roche), 1 mM serine protease inhibitor PMSF (phenylmethylsulfonyl fluoride) and 0.5 mM DTT for 10 min at 4 °C and centrifuged at 16400 rpm (25 000 g) for 10 min at 4 °C. Cells lysed with Laemmli buffer were suspended in an appropriate amount of 3 × Laemmli buffer and boiled for 5-10 min. The protein concentration of the lysates in lysis buffer was determined using the Bradford method.

Cell lysates were resolved on an 8% sodium dodecyl sulfate-polyacrylamide gel (SDS-PAGE) and transferred to a 0.45 µm pore size nitrocellulose membrane (Protran). The membranes were boiled for 10 min in ultrapure water H_2_O directly after transfer and blocked with 5% skimmed milk powder in 0.3% PBS-Tween20. The primary antibodies were diluted in PBS containing BSA and 0.02% NaN_3_. The membranes were incubated with the primary antibodies overnight at 4 °C. The primary antibodies used were: anti-HSF1 (ADI-SPA-901, Enzo Life Sciences), anti-HSF2 (clone 3E2, MAB88079, Sigma-Aldrich or HPA031455, Sigma-Aldrich), anti-HSP70 (ADI-SPA-810, Enzo Life Sciences), anti-FLAG (clone M2, F3165 M2ab, Sigma-Aldrich), anti-Myc (clone 9E10, M5546, Sigma-Aldrich) and anti-β-tubulin (clone AA2, T8328, Sigma-Aldrich). Secondary antibodies conjugated to horseradish peroxidase were purchased from Promega, GE Healthcare, or Abcam.

### Biotin-mediated oligonucleotide pulldown assay

The biotin-mediated oligonucleotide pulldown assay is modified from (Anckar et al. 2006). U2OS WT cells were lysed with buffer C (25% glycerol, 0.42 M NaCl, 1.5 mM MgCl_2_, 0.2 mM EDTA, pH 8.0, 20 mM HEPES) containing 0.5 mM DTT and 0.5-1 mM PMSF. Buffer C extracts (150 – 200 µg protein) were incubated in binding buffer (20 mM Tris-HCl, pH 7.5, 100 mM NaCl, 2 mM EDTA, 10% glycerol) with 3 µg annealed oligonucleotides, containing either a heat shock element (HSE) or a scrambled sequence, and 0.5 µg/µl salmon sperm DNA (Sigma-Aldrich). The HSE-containing oligonucleotides 5’-biotin-TCGACTAGAAGCTTCTAGAAGCTTCTAG-3’ and 5’-CTAGAAGCTTCTAGAAGCTTCTAGTCGA-3’ (Vuori et al. 2009), and the scrambled control oligonucleotides 5’-biotin-AACGACGGTCGCTCCGCCTGGCT-3’ and 5’-AGCCAGGCGGAGCGACCGTCGTT-3’ (Anckar et al. 2006) were purchased from Oligomer. The proteins were allowed to bind to the oligonucleotides for 20 min at room temperature and 30 min at 4 °C. The samples were precleared with Glutathione Sepharose 4 Fast Flow (17-5132-01, GE Healthcare) for 30 min at 4 °C under rotation. The remaining DNA was precipitated with 25 µl Streptavidin-Sepharose 4B (434341, Invitrogen) for 1 h at 4 °C under rotation. Bound fractions were washed three times with binding buffer and twice with binding buffer containing 0.2% Triton X-100. The DNA-bound proteins were suspended in 20 µl 3x Laemmli buffer and boiled for 5 min to elute the proteins. The samples were analyzed with SDS-PAGE and immunoblotting.

### Quantitative real-time reverse transcription-PCR

RNA was isolated from U2OS and HeLa cell pellets using a Nucleospin RNA isolation kit (Macherey-Nagel) according to the manufacturer’s instructions and the RNA concentration was measured using a NanoDrop 2000 spectrophotometer (Thermo Fisher Scientific). The reverse transcriptase enzyme M-MLV RT (H-) (Moloney Murine Leukemia Virus Reverse Transcriptase RNase H minus, Promega) was then used to transcribe 1 µg of total RNA to cDNA using Oligo(dT)_15_ primers (Promega). KAPA Probe Fast qPCR Master Mix (2X) ABI Prism (for *hsp70, hsp110* and *18S* RNA) or a KAPA SYBR FAST qPCR Master Mix (2X) ABI Prism (for *satIII* and *hGAPDH*) kits (KK4706 and KK4604, KapaBiosystems) were used for the qRT-PCR reactions and these were performed with a StepOnePlus Real-Time PCR system (Applied Biosystems). Primers and probes were purchased from Oligomer and Roche Universal Probe Library, and can be found in Table 2. The relative quantities of *hsp70* and *hsp110* were normalized against *18S* rRNA with the help of a standard curve. The relative quantities of *satIII* were normalized against *hGAPDH*. The fold induction was calculated against the respective mRNA levels in control cells. All reactions were run in triplicate from samples, generally derived from at least three biological replicates.

**Table 2.**
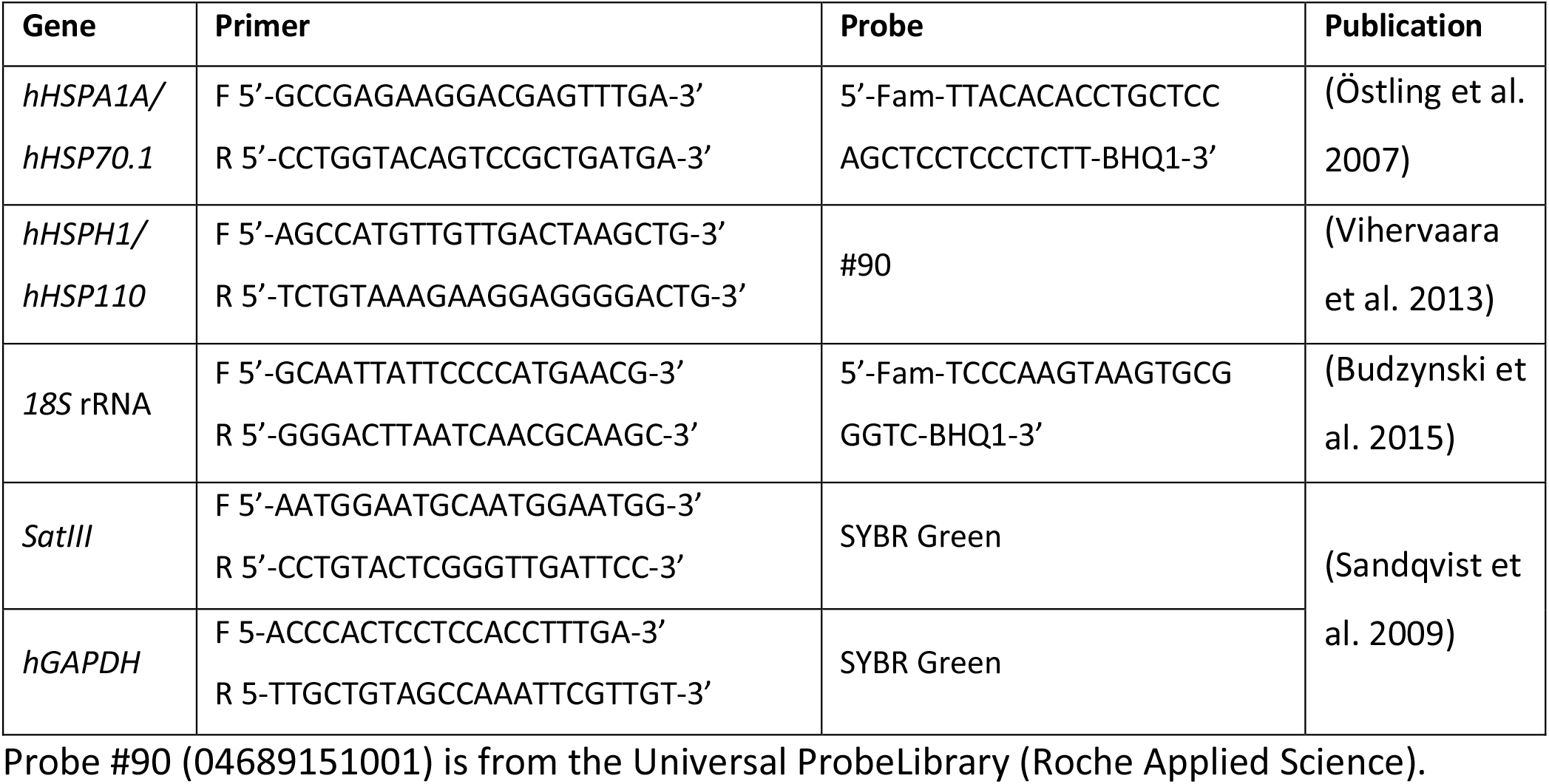
Primers (F: forward, R: reverse) and probes used for the qPCR reactions

### Statistical analysis

Statistical analyses were performed using GraphPad Prism 7 Software (GraphPad Prism Software, http://www.graphpad.com). The data was analyzed using one-way ANOVA and corrected with the Holm-Šídák’s multiple comparisons test. The significance level was set to 0.05. Mean + SEM is shown in the figures.

### Immunofluorescence

To detect nuclear stress bodies, immunofluorescence was performed as in (Sandqvist et al. 2009). Briefly, HeLa cells were cultured on coverslips and fixed with 100% methanol for 6 min at 4 °C. The methanol was aspirated and the cells were washed three times with 0.05% PBS-Tween20. The cells were incubated in a blocking solution containing 10% BSA (bovine serum albumin) in 0.05% PBS-Tween20 for 1 h. The cells were incubated with anti-HSF1 antibody (Holmberg et al. 2000) diluted in blocking solution (1:300) overnight at 4 °C. The secondary antibody (anti-rabbit IgG, Alexa Fluor 488) was diluted in blocking solution (1:700) and added for 1 h. The coverslips were mounted using VECTASHIELD mounting medium (Vector Laboratories) and the cells were visualized with an LSM510-Meta scanning confocal microscope (Carl Zeiss).

### HSP90-HSF1 interaction assay

HSP90-HSF1 interaction was studied as in (Kijima et al. 2018). HEK293 cells were transfected with WT HSF1 and either of the two HSP90 mutants: FLAG-HSP90α E47A and FLAG-HSP90β E42A. Both mutant constructs were a kind gift from Dr. Len Neckers (NIH, Bethesda, USA). These mutants are in a closed conformation and are described in detail in (Kijima et al. 2018). Transfections were performed using the Neon Transfection System (Thermo Fisher Scientific) according to the manufacturer’s instructions. 7 × 10^6^ HEK293 cells were suspended in 100 µl Resuspension Buffer R and mixed with plasmids for Myc-His-HSF1 WT (described in (Westerheide et al. 2009)), pBlueScript empty vector (BS), FLAG-HSP90α E47A or FLAG-HSP90β E42A. The BS vector was used as a negative control. The cells were subjected to electroporation (1245 V, 10 ms pulse width, 3 pulses), plated and left to recover in culture medium for 48 h before treatments.

For immunoprecipitation, the cells were lysed with TGNET buffer (50 mM Tris HCl pH 7.5, 5% Glycerol, 100 mM NaCl, 2 mM EDTA, 0.5% Triton X-100) containing 1 mM PMSF, and protease and phosphatase inhibitor cocktails (04693159001 and 04906845001, Roche). Lysates were incubated on ice for 10 min and centrifuged at 16400 rpm (25 000 g) for 15 min at 4 °C. The protein concentrations of the supernatants were determined using the Bradford method and 15 µg of protein was subjected to SDS-PAGE for protein expression analyses. 700 µg of protein was subjected to immunoprecipitation using 30 µl of Anti-FLAG M2 affinity beads (A2220, Sigma-Aldrich). The beads were incubated with rotation for 2 h, at 4 °C and then centrifuged at 6000 rpm (3500 g) for 30 s. The beads were washed three times with TGNET buffer and then suspended in 12 µl 3 × Laemmli buffer and boiled for 5 min to elute the proteins. The samples were analyzed with SDS-PAGE and immunoblotting.

## Results

### Gambogic acid (GB) induces a heat shock response in multiple cell lines

GB has been extensively studied in the context of cancer, and shown to work as an anticancer agent (Banik et al. 2018). However, there are contradicting reports whether GB induces a HSR (Davenport et al. 2011; Yim et al. 2016), which is cytoprotective for cancer cells and therefore a potentially problematic feature for an anticancer drug. Therefore, to determine if GB indeed induces a HSR, we treated both transformed (U2OS and HeLa), and untransformed (HDF and RWPE-1) cells with acute treatments of GB (Fig. 1a). HSF1 hyperphosphorylation and upregulation of HSP70 protein levels were used as proxies for the activation of the HSR (Sarge et al. 1993). We included the previously defined HSP90-inhibitors celastrol and 17-AAG (17-allyllaminogeldanamycin) in the analyses, since both are known activators of HSF1 and subsequent HSR (Bagatell et al. 2000; Westerheide et al. 2004). As a positive control for HSF1 hyperphosphorylation we used 1 h heat shock (HS) treatment at 42 °C and for HSP70 protein upregulation, HS + 3 h recovery (H+R) was used. In both untransformed and transformed cell lines, acute treatments with GB induced HSF1 hyperphosphorylation and increased HSP70 levels to a similar extent as HS (Fig. 1a). Moreover, HSF1 hyperphosphorylation and HSP70 upregulation were also induced by celastrol in all cell lines, whereas 17-AAG at these concentrations induced an HSR in all cell lines except primary HDFs. Altogether, these results demonstrate that GB induces a robust HSR in untransformed and transformed human cells.

**Fig. 1.**
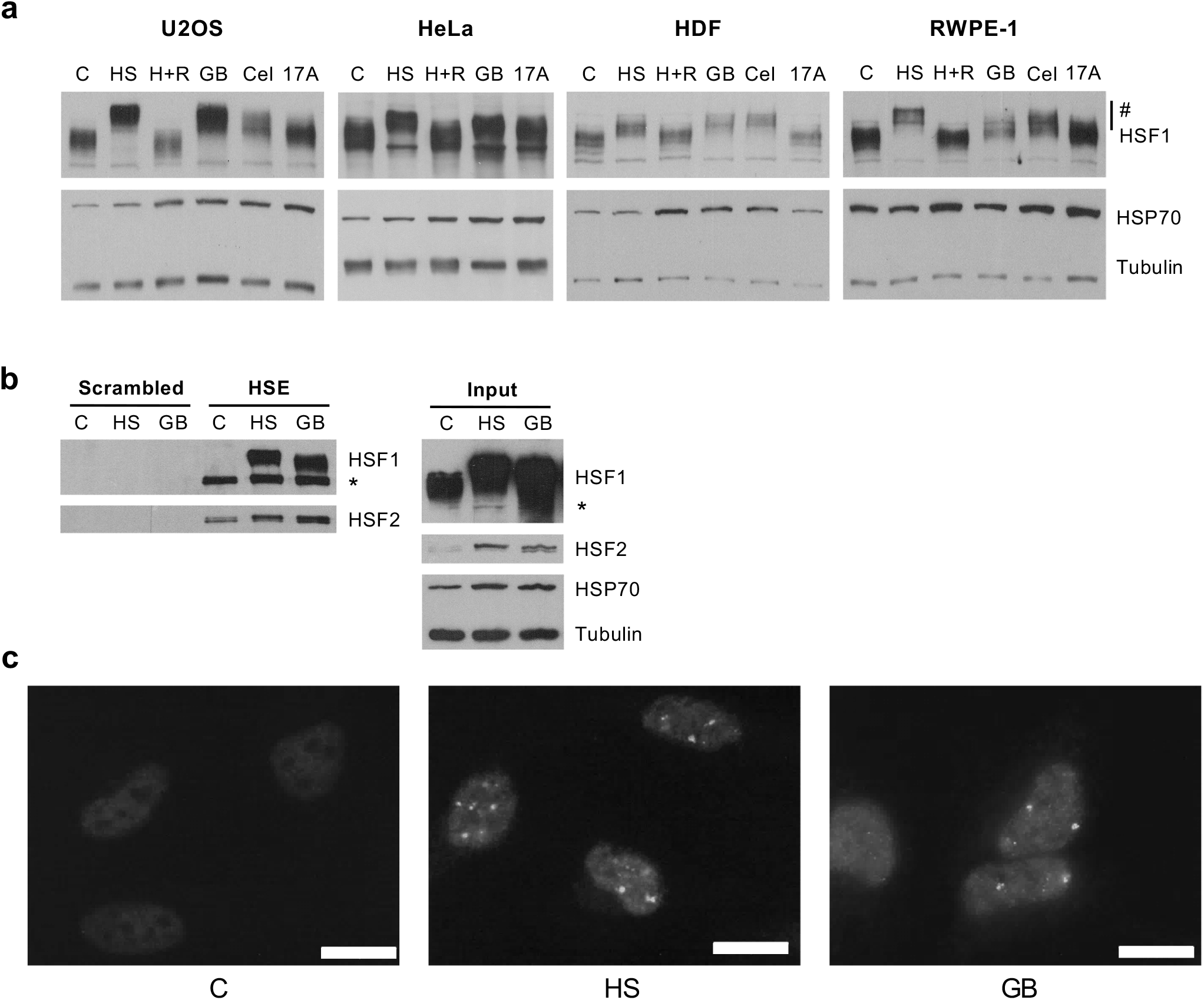
Acute treatment with gambogic acid induces a heat shock response (HSR). **a** Immunoblot analysis of HSF1 and HSP70 expression. U2OS, HeLa, HDF and RWPE-1 cells were treated with heat shock (HS, 42 °C, 1 h), 1 h HS with 3 h recovery at 37 °C (H+R), and gambogic acid (GB), celastrol (Cel) or 17-AAG (17A) for 4 h. U2OS: 1.25 μM GB and Cel, 0.25 μM 17A; HeLa: 2 μM GB, 1.5 μM 17A; RWPE-1: 1.25 μM GB and Cel, 3 μM 17A; HDF: 2.5 μM GB and Cel, 3 μM 17A. The HSF1 uppershift induced by HS and treatments (labeled with #) correspond to hyperphosphorylated HSF1 forms. Tubulin was used as a loading control. **b** Oligonucleotide-mediated pulldown of HSF1 and HSF2 in WT U2OS cells, untreated (C), treated with heat shock 42 °C for 1 h (HS), or with 1.25 μM gambogic acid for 4.5 h (GB). Input indicates total cell lysates from treated cells. Tubulin serves as a loading control. * indicates previously blotted HSF2. **c** Immunofluorescence staining of endogenous HSF1 in nuclear stress bodies in HeLa cells. GB: 4 µM GB, HS: 2 h heat shock (HS, 42°C) (n=3). Scale bar: 20 µm

To further address whether GB treatment results in increased HSF activity, we employed an oligopulldown assay to study both HSF1 and HSF2 DNA-binding capacity (Fig. 1b). Lysates from treated cells were subjected to a biotinylated heat shock element (HSE)-containing oligo or a scrambled oligo, which were purified with streptavidin beads together with their respective binding proteins. HSF2 bound to the HSE-oligo already in untreated cells, and the binding was increased after treatments with GB or HS (Fig. 1b), in agreement with previous results in heat-shocked cells using chromatin immunoprecipitation (Ahlskog et al. 2010). In contrast, HSF1 DNA-binding was detected only after HS or GB-treatment. Therefore, acute treatment with GB impacts the DNA-binding activities of both HSF1 and HSF2, in a manner similar to heat shock.

HSF1 and HSF2 localize to subnuclear structures called nuclear stress bodies (nSBs) upon heat stress (Alastalo et al. 2003; Jolly et al. 2002; Jolly et al. 2004). These nSBs form on areas with pericentromeric heterochromatin, where HSF1 induces transcription of noncoding *satellite III* (*satIII*) RNA. The function of the non-coding RNAs is not well understood, but the transcripts have been suggested to affect chromatin organization and recruitment of transcription and splicing factors (Biamonti and Vourc’h 2010), and HSF1 recruitment to the nSBs is a hallmark of the HSR in human cells. Using indirect immunofluorescence, probing for endogenous HSF1, we studied the localization of HSF1 after HS and GB treatment. We demonstrated that GB induces HSF1-localization to nSBs in HeLa cells, similarly to HS (Fig. 1c). Taken together, our results show that the HSR induced by acute treatment with GB activates both HSF1 and HSF2 in multiple human cell lines of different origin.

### GB and GBA induce a heat shock response in a thiol-dependent manner

The natural products, celastrol and GB, have similar chemical features, as they both contain an α,β-unsaturated ketone moiety (Fig. 2a). The resin from *Garcinia hanburyi* contains an additional active compound called gambogenic acid (GBA; Fig. 2a), which also contains an α,β -unsaturated ketone moiety. The biological effects of celastrol can be inhibited by the excess of free thiol, suggesting that celastrol reacts with key thiols in proteins (Lee, J. H. et al. 2006; Lee, J. Y. et al. 2015; Peng et al. 2010; Trott et al. 2008). To examine whether GB is also thiol-responsive and inactivated by the excess of free thiols, we incubated GB with 10-fold excess dithiothreitol (DTT) before applying it to the cells. Intriguingly, the results showed that GB ability to induce *HSP* gene expression was inactivated by DTT, as evidenced by significantly lower *HSPA1A* (*HSP70*) mRNA expression levels in cells treated with a GB+DTT mixture (Fig. 2b). Moreover, extensive HSF1 hyperphosphorylation was also abolished after GB+DTT treatment (Fig. 2c). We also determined the amount of *satIII* transcripts produced after treatment with GB. In agreement with nSB formation (Fig. 1c), we observed that the transcription of *satIII* RNA was induced by GB, as well as HS (Fig. 2d). The GB-induced *satIII* transcripts were not produced if GB was pretreated with DTT before addition to the cells (Fig. 2d, e), demonstrating that GB can indeed be inhibited by the excess of thiols. We also addressed whether GBA can elicit a HSR. We found that acute GBA treatments also induced a HSR and that GBA was inactivated by incubation with excess DTT, suggesting that both GB and GBA act in a thiol-dependent manner on the triggering of a HSR (Fig. 2f).

**Fig. 2.**
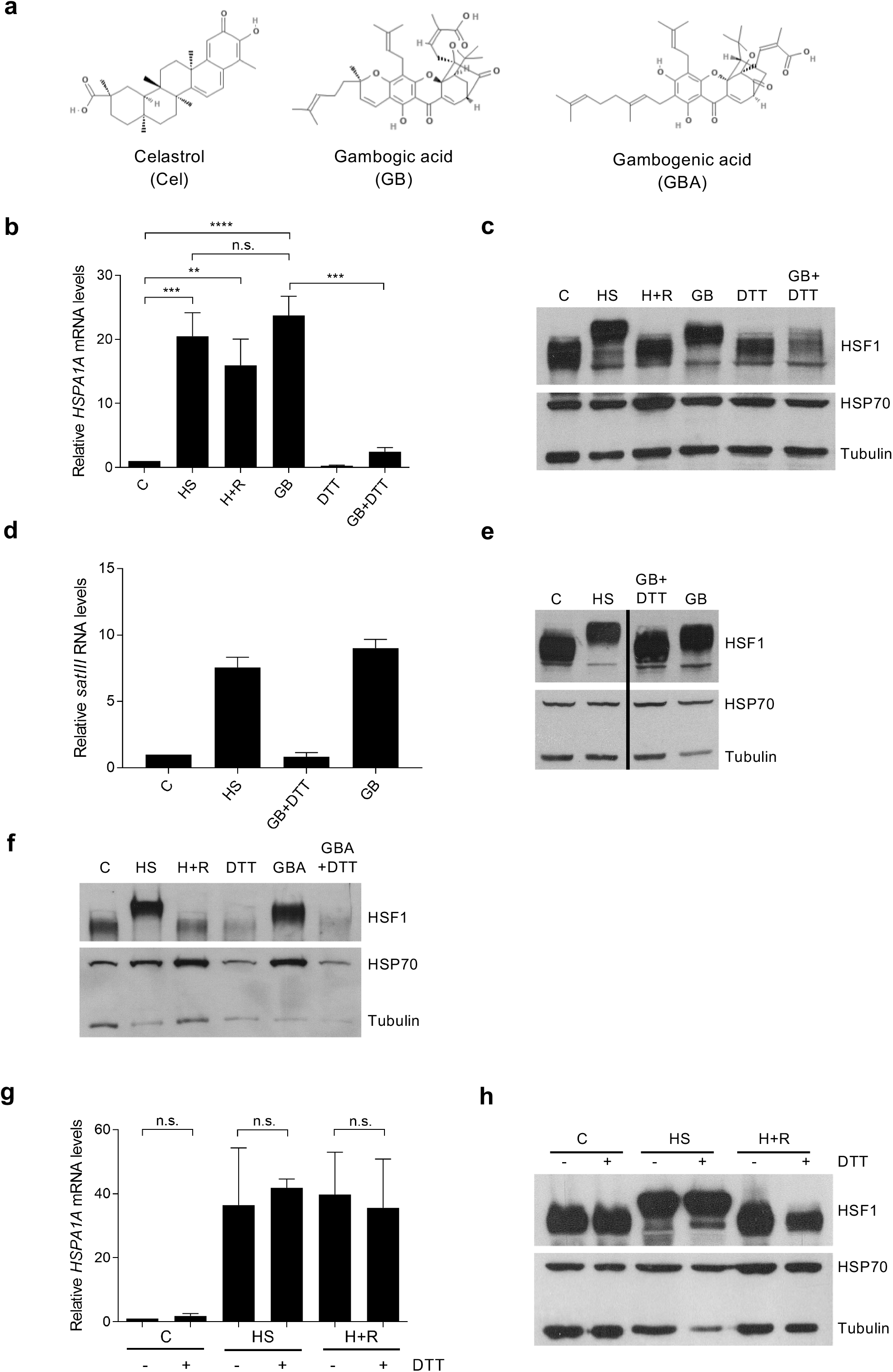
Gambogic acid and gambogenic acid induce a HSR in a thiol-dependent manner. **a** Molecular structures of celastrol, gambogic acid (GB) and gambogenic acid (GBA). The structures are taken from PubChem (https://pubchem.ncbi.nlm.nih.gov, PubChem IDs: 122724, 15559465 and 10794070) and modified. **b** qRT-PCR of *HSPA1A (HSP 70)* mRNA of WT U2OS cells treated with 1 h heat shock at 42 °C (HS), or with 1 h HS followed by 3 h recovery at 37 °C (H+R), 1.25 µM GB for 4.5 h or with GB pretreated with 10xDTT for 15 min prior to adding the mixture to the cells (GB+DTT) or with 12.5 µM DTT alone for 4.5 h. *HSPA1A (HSP70)* mRNA normalized to 18S rRNA (n=3). One-way ANOVA, mean ± SEM shown. ** p≤ 0.01, *** p≤0.001, ****p≤0.0001, n.s. p >0.05. **c** Immunoblot analysis of HSF1 and HSP70 of the corresponding samples in **b**. Tubulin was used as a loading control. **d** qRT-PCR of *satIII* transcripts in HeLa cells treated with 1 h HS (42 °C) or for 4 h with 4 µM GB or 4 µM GB pretreated with 10xDTT (GB+DTT). qRT-PCR with SYBR Green, normalized to *hGAPDH* (n=2). Mean ± SEM shown. **e** Immunoblot analysis of HSF1 and HSP70 expression of the corresponding samples in **d**. Tubulin was used as a loading control. **f** Immunoblot analysis of U2OS cells treated with 1 h heat shock at 42 °C (HS), or with 1 h HS followed by 3 h recovery at 37 °C (H+R), 2.5 µM GBA for 4.5 h or with GBA pretreated with 10xDTT for 15 min prior to adding the mixture to the cells (GB+DTT) or with 25 µM DTT alone for 4.5 h. **g** qRT-PCR of *HSPA1A* mRNA from WT U2OS cells that were treated with heat shock (HS, 42 °C, 1 h), 1 h HS with 3 h recovery at 37 °C (H+R), with our without 12.5 µM DTT (-/+DTT) prior to treatment. C+DTT samples were treated with 12.5 µM DTT for 1 h. *HSPA1A* mRNA normalized to *18S* rRNA (n=3). One-way ANOVA, mean ± SEM shown. n.s. p >0.05. **h** Immunoblot analysis of HSF1, HSF2 and HSP70 of the corresponding samples in **g**. Tubulin was used as a loading control

Huang and coworkers (1994) showed that the HSR can be inhibited by thiol reducing agents (e.g. 2 mM DTT) (Huang, L. E. et al. 1994). To rule out that DTT does not, by itself, inhibit the HSR in our experiments, we treated cells before HS treatment with the same concentrations of DTT as in Fig. 2a and b (12.5 µM), and assessed HSP70 mRNA and protein levels, as well as HSF1 hyperphosphorylation. We observed normal HSR and recovery profiles, showing that 12.5 µM DTT does not inactivate the HSR by itself (Fig. 2g, h). Therefore, we conclude that GB and GBA, as celastrol, are thiol-responsive chemicals and that pretreatment with DTT inactivates GB and GBA and therefore perturbs the GB/GBA-induced activation of the HSR.

### The GB- or GBA-induced HSR is HSF1-dependent

To study whether HSF1 and HSF2 are required for GB- or GBA-induced HSR, we treated WT U2OS cells as well as CRISPR-generated *HSF1* or *HSF2* knockout (KO) U2OS cells with GB and GBA. 17-AAG was used as positive control, because it has been shown to induce a HSF1-dependent HSR (Bagatell et al. 2000). Using immunoblotting, we show that all compounds induced a robust HSF1 hyperphosphorylation and upregulation in HSP70 protein expression levels in WT cells (Fig. 3a). No induction of HSP70 protein was detected with any of the treatments in *HSF1* KO cells (Fig. 3a). In cells lacking HSF2, HSF1 was robustly hyperphosphorylated in response to GB and GBA, but less in response to 17-AAG. There seems to be slightly less HSP70 protein in cells lacking HSF2.

**Fig. 3.**
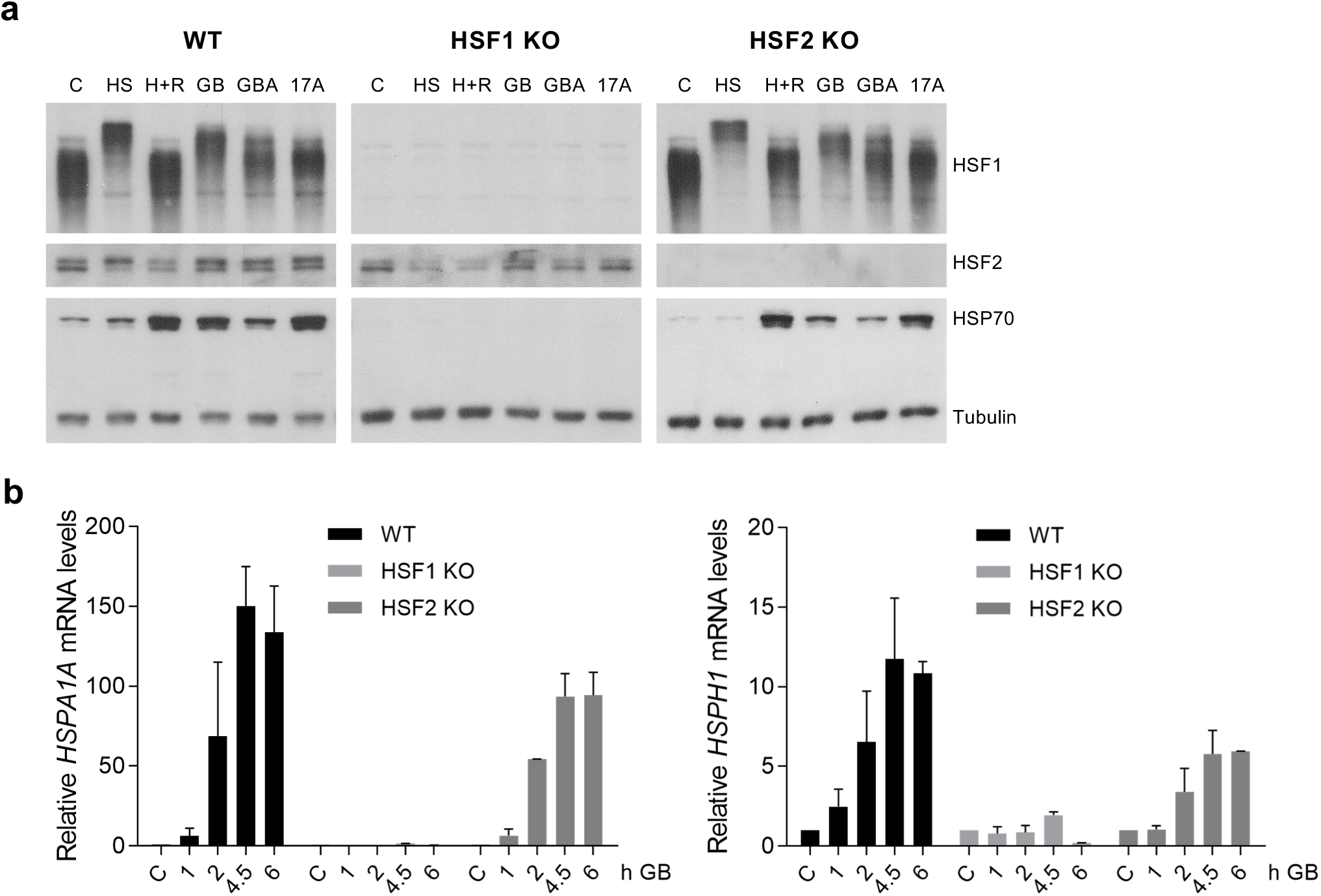
Gambogic acid and gambogenic acid induce a heat shock response (HSR) in an HSF1-dependent manner, whereas HSF2 is dispensable for the HSR. **a** WT, *HSF1* and *HSF2* knockout (KO) U2OS cells were treated with heat shock (HS, 42 °C, 1 h), 1 h HS with 3 h recovery (H+R), 1.25 μM gambogic acid (GB), 2.5 μM gambogenic acid (GBA) or 0.5 μM 17-AAG for 4.5 h. Immunoblot of HSF1, HSF2 and HSP70. Tubulin serves as a loading control. **b** WT, *HSF1* KO and *HSF2* KO U2OS cells were treated with 1.25 μM GB for indicated times. qRT-PCR of *HSPA1A (HSP70)* and *HSPH1 (HSP110)* mRNA, normalized to *18S* rRNA (n=2). Mean ± SEM shown

The effect of GB on mRNA levels of *HSPA1A* (*HSP70*) and *HSPH1* (*HSP110*) at different time points in WT, *HSF1* KO, and *HSF2* KO U2OS cells was investigated using quantitative RT-PCR. As demonstrated in Fig. 3b, both *HSPA1A* and *HSPH1* mRNA levels increased time-dependently after treatment with GB both in WT and in *HSF2* KO cells, albeit the levels of *HSPA1A* and *HSPH1* mRNA were lower in *HSF2* KO cells. There was no induction of *HSPA1A* or *HSPH1* mRNA in response to GB treatment in cells lacking HSF1.

These results show that 17-AAG, GB and the GB-analogue GBA induce a HSR that is strictly dependent on HSF1 but not on HSF2. In accordance with previous studies on HS-induced HSR (Joutsen et al. 2020; Östling et al. 2007; Vihervaara et al. 2013), HSF2 indeed modulates the HSR induced by GB and the other compounds.

### GB and GBA disrupt the interaction between HSP90 and HSF1 or HSF2

GB has been proposed to be an HSP90 inhibitor (Davenport et al. 2011; Yim et al. 2016). The proposed model of action for HSP90 inhibitors inducing a HSR is that they disrupt HSP90-HSF1 interaction, thereby freeing HSF1 that can be activated (Åkerfelt et al. 2010). However, the HSP90-HSF1 interaction is transient and weak (Zou et al. 1998), and it is challenging to co-immunoprecipitate HSF1 with HSP90 without crosslinking. HSP90 functions as a dimer and cycles between a closed ATP-bound state and an open state where ATP is hydrolyzed or absent (Pearl 2016). By mutating glutamic acid residues to alanines at positions 47 in HSP90α and 42 in HSP90β (E47A and E42A, respectively), Kijima and coworkers (2018) generated two HSP90 mutants that are constantly in the closed ATP-bound conformation (Fig. 4a). Using immunoprecipitation without crosslinking they showed that these closed conformation HSP90 mutants can stably bind to HSF1, and that N-terminal HSP90 inhibitors, like 17-AAG, disrupt the HSP90-HSF1 interaction (Kijima et al. 2018).

**Fig. 4.**
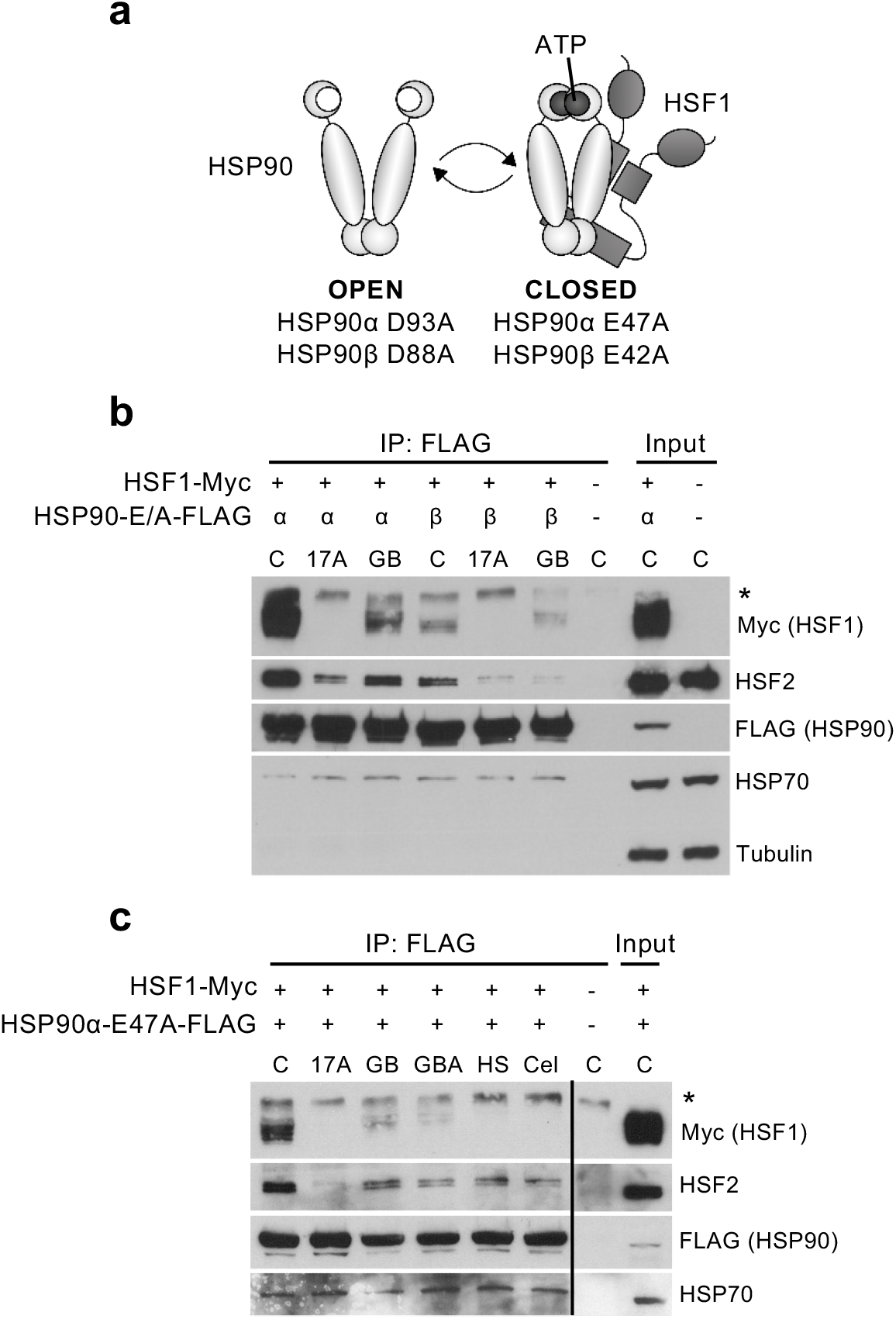
Gambogic acid and gambogenic acid induce a heat shock response by disrupting HSP90-HSF1/HSF2 interaction. **a** Schematic model of the HSP90-mutants. The closed form mutants can interact with HSF1, whereas the open mutants cannot (Kijima et al., 2018). **b** HEK293 cells were transfected with either FLAG-HSP90α E47A (α) or FLAG-HSP90β E42A (β) (closed form mutants) and HSF1-Myc, and treated two days later with 10 μM 17-AAG (17A) or 1.5 μM gambogic acid (GB) for 4 h. BlueScript empty vector (-) was transfected as a negative control. HSP90-FLAG was precipitated with FLAG-beads and exogenous HSF1 (Myc), HSF2, exogenous HSP90 (FLAG) and HSP70 were analyzed by immunoblotting. Tubulin was used as a loading control in the whole cell lysates (WCL). * indicates an unspecific band. **c** HEK293 cells were transfected with FLAG-HSP90α E47A and HSF1-Myc and treated two days later for 4h with 10 μM 17-AAG (17A), 1.6 μM gambogic acid (GB), 3.2 μM gambogenic acid (GBA), 3 μM celastrol (Cel) or heat shocked (3 h, 42 °C). BlueScript empty vector (-) was transfected as a negative control. * indicates an unspecific band. The samples were analyzed as in **b**

There are conflicting reports regarding GB binding to HSP90 (Davenport et al. 2011; Yim et al. 2016), and GBA has to the best of our knowledge not been studied in the context of HSP90. Here, we investigated if GB and GBA also disrupt the interaction between HSF1 and HSP90α and β closed form mutants. We co-transfected HEK293 cells with WT HSF1-Myc-His and FLAG-HSP90α E47A or FLAG-HSP90β E42A. In accordance with Kijima and coworkers (Kijima et al. 2018), HSP90α E47A forms a stronger interaction with HSF1 than HSP90β E42A as more HSF1 (Myc) was immunoprecipitated with HSP90α E47A than with HSP90β E42A (Fig. 4b). 17-AAG completely disrupted the interaction between HSF1 and HSP90α E47A and HSP90β E42A. We observed that GB also disrupts the interaction between HSF1 and HSP90, both α and β, albeit with lower efficiency than 17-AAG (Fig. 4b).

In addition to its extremely large repertoire of clients, including multiple oncogenic kinases and key transcription factors, HSP90 also interacts with other components of the protein folding machinery, such as HSP70. As seen in Fig. 4b, endogenous HSP70 was immunoprecipitated with both HSP90α E47A and HSP90β E42A but this interaction was not affected by 17-AAG or GB. Interestingly, we also observed that endogenous HSF2 was immunoprecipitated by FLAG-tagged HSP90α and β (Fig. 4b). To our knowledge, this is the first report demonstrating an HSP90-HSF2 interaction. Both 17-AAG and GB disrupted the interaction between HSF2 and HSP90α or β but to a different degree than HSF1 (Fig. 4b), suggesting that HSF1 and HSF2 can bind independently to HSP90.

The interaction between HSP90α E47A and HSF1 was more robust and hence we explored if other compounds known to induce a HSR would disrupt the interaction between HSP90α E47A and HSF1. 17-AAG, as well as HS and celastrol, completely disrupted the interaction between HSF1 and HSP90α (Fig. 4c). GB and GBA also disrupted the interaction, with slightly lower efficacy. HSF2 interaction with HSP90 was disrupted by all treatments, with 17-AAG treatment causing the most effective disruption. Endogenous HSP70 interaction with HSP90α E47A was not disrupted by these different compounds.

Taken together, we show that GB and GBA treatment can disrupt the interaction between HSP90 and HSF1. We propose that this may be part of the mechanism by which GB and GBA activate the HSR. We also show that endogenous HSF2 and HSP70 are found in the HSP90-HSF1 complex.

## Discussion

Targeting HSP90 has been considered beneficial in cancer treatment with HSP90 having many so-called client proteins that are required for a rapidly growing cell. Gambogic acid (GB), a bioactive natural product and potential HSP90-inhibitor, has been shown to kill cancer cells more readily than normal cells, which would support GB as an anticancer drug. However, the molecular mechanisms of GB are still not clear. Here, we show that acute treatments with either GB or its structural analog GBA induce a thiol-dependent HSR in multiple cell lines derived from different cellular origins. We demonstrate that GB and GBA treatment induces a HSR, at least partially by disrupting the HSP90-HSF1 and HSF2 interaction.

Unfortunately, most HSP90 inhibitors tested today induce an HSR, which is cytoprotective during cancer treatments, as higher levels of HSPs are favorable for the cancer cell (Neckers and Workman 2012). Interestingly, a recent study shows that using HSP90 inhibitors at continuous low, non-HSR inducing levels improves immunomodulatory functions and is a promising step in immunotherapeutic strategies for treating cancer (Jaeger et al. 2019). Therefore, further studies are required to test if already developed HSP90 inhibitors could be used at lower, non-HSR inducing levels, and perhaps as combinatorial therapies.

Molecules containing α,β-unsaturated ketone moieties are highly reactive and covalently modify a plethora of functional cysteine residues in many proteins (Weerapana et al. 2008). Thiol-reactive compounds have previously been shown to activate HSF1 in yeast and to induce a HSR in mammalian cells (Dayalan Naidu and Dinkova-Kostova 2017; Santagata et al. 2012; Trott et al. 2008; Wang et al. 2012). GB and GBA contain an α,β-unsaturated ketone moiety and reducing this moiety at carbons C9-C10 renders GB inactive (Han et al. 2005). We show that the HSR, which is induced by GB or GBA, is inhibited in the case GB or GBA is incubated with excess free thiols (Fig. 2). Our observations are corroborated by previous studies showing that GB can bind covalently to cysteine residues of different proteins, a process that is inhibited by excess free thiol such as DTT and thiol-containing antioxidants (Palempalli et al. 2009; Seo et al. 2019; Yang et al. 2012). Importantly, our study provides the first demonstration that GBA induces a HSR and is thiol-responsive.

We also bring new insights in to the mechanisms of action of GB and GBA. N-terminal, but not C-terminal HSP90 inhibitors disrupt the interaction between HSF1 and closed-form HSP90 mutants (Kijima et al. 2018). Here we show that both GB and GBA treatments can disrupt the interaction between HSP90, HSF1 and HSF2, suggesting that GB and GBA are mechanistically similar to N-terminal inhibitors. Davenport and coworkers reported, using surface plasmon resonance, that GB binds to the N-terminal part of HSP90, whereas Yim and coworkers reported that biotinylated GB binds to the middle domain of HSP90 (Davenport et al. 2011; Yim et al. 2016). So far, studies have shown that normal cells lines are less sensitive to GB than cancer cell lines, perhaps due to redox homeostasis (Duan et al. 2014). However, it will be challenging to use GB and GBA as specific cancer drugs due to the thiol-reactive nature of GB and GBA, as they may interact with many proteins in the cell. Alternatively, it may be that GB and GBA can be used at lower doses in combinatorial therapies. GB has been shown to enhance the cytotoxic effects of different chemotherapeutic agents, e.g. adriamycin, docetaxel and oxaliplatin in different cancer cells (Banik et al. 2018). Since GB has a poor aqueous solubility, poor biodistribution and multi target capacity, nanotechnology approaches are currently being developed (Hatami et al. 2020). It also has to be determined whether GB and GBA could be used as treatments for diseases where induction of a HSR and an increase of HSPs is beneficial. For example, celastrol, which inhibits HSP90 and also contains an α,β-unsaturated ketone moiety has cardioprotective effects in rat ischemia/reperfusion studies via induction of HSPs and heme oxygenase 1 (HO-1) (Aceros et al. 2019), and also neuroprotective effects (Cascao et al. 2017).

We also bring novelty by showing that HSF2 participates in a complex comprising HSF1 and HSP90 and in regulation of HSR by GB and GBA. We show that GB- and GBA-induced HSR is HSF1-dependent and modulated by HSF2. Previous studies have showed that HSF2 also modulates the HS-induced HSR (Joutsen et al. 2020; Östling et al. 2007; Vihervaara et al. 2013). HSF2 is a short-lived protein and its activity is mainly regulated by its cellular expression levels and post-translational modifications (Ahlskog et al. 2010; Björk and Sistonen 2010; Mathew et al. 1998; Park et al. 2015). Both HSF1 and HSF2 are implicated in cancer and HSF2 is downregulated in malignant cancers (Björk et al. 2016; Dai 2018; Puustinen and Sistonen 2020). HSF2 functions as a suppressor of prostate cancer invasion (Björk et al. 2016). To our knowledge, the role of HSF2 in response to HSP90 inhibitors has not previously been thoroughly studied. Interestingly, a recent study showed that *HSF2* KO cells are more sensitive to HSP90 inhibitors, therefore, further studies regarding the role of HSF2 in the context of HSP90 inhibition is of importance (Joutsen et al. 2020).

Studying the interaction between endogenous HSP90 and HSF1 is challenging, reflecting a possible transient interaction between the two proteins. We show that both HSF1 and HSF2 are found in a complex with HSP90 closed form mutants. Kijima and coworkers mapped the interaction between closed form HSP90 mutants and HSF1, and determined that both HSF1 heptad repeats HR-A/B and a part of the regulatory RD domain are required for HSP90 interaction (Kijima et al. 2018). HSF2 also contains heptad repeats required for trimerization with HSF1, whereas the regulatory domain of HSF2 does not contain multiple phosphorylation sites (Gomez-Pastor et al. 2018). From our results we cannot determine if HSF2 binds directly to HSP90 as a potential homotrimer, or if the interaction is via its trimerization with HSF1. It is also unclear if HSF2 complex formation with HSP90 can regulate HSF2 stability. Therefore, it is important to include HSF2 in further studies regarding HSP90 and HSF1. HSP90 has long been assumed to participate in keeping HSF1 monomers from trimerizing in the cytosol but has recently been shown to bind to HSF1 trimers and remove them from HSEs (Kijima et al. 2018). HSP90β also takes part in the activation of HSF1 by lowering the temperature required for heat-induced HSF1 trimerization (Hentze et al. 2016). It is likely that HSP90 participates in the regulation of HSR in a more prevalent and complex manner than previously anticipated, to which HSF2 adds an additional exciting and novel aspect.

In conclusion, we show that the natural products GB and GBA, isolated from gamboge resin, induce a strong HSR, in a thiol-dependent manner, and leads to the release of HSF1 and HSF2 from the HSP90-HSF complex. The GB and GBA-dependent activation of the HSF-pathway should be taken into account when using these compounds in different studies. In particular, this is a limiting step in using them in anti-cancer therapies, however, there may be other pathologies, where an activation of the HSR by GB or GBA would be beneficial, through for example neuroprotective effects.

## Authors’ contributions (optional: please review the submission guidelines from the journal whether statements are mandatory)

LP, SS and PRM designed the research

AdT, VM, and DSD generated and analyzed the U2OS cell lines

LP, SS, VB and PRM performed the experiments

LP, SS and PRM analyzed the data

LP and PRM wrote the manuscript and all authors made editorial contributions to the final manuscript

## Acknowledgments

Acknowledgments of people, grants, funds, etc. should be placed in a separate section on the title page. The names of funding organizations should be written in full.

We thank the laboratories of Keith Joung (Massachusetts General Hospital) and George Church (Harvard University) for reagents for generating the CRISPR cell lines. We thank Jean-Paul Concordet (MNHN, Paris) for guidance with the CRISPR cell lines. Ingrid Karppi is acknowledged for her assistance with GBA experiments while in Roos-Mattjus lab. We thank Lea Sistonen, Jenny Joutsen and the members of the Sistonen laboratory for their valuable comments and critical review of the manuscript; members of the Mattjus laboratory for technical support; and J Peter Slotte for all the support as the department head when this study was initiated.

This study has been funded by Åbo Akademi University Research Foundation (PRM), Magnus Ehrnrooth Foundation (PRM), Swedish Cultural Foundation in Finland (PRM) and Medical Research Foundation Liv och Hälsa (PRM). VM was funded by the CNRS (Projet International de Coopération Scientifique PICS 2013-2015) for her collaboration with PRM and by the Short Researcher Mobility France Embassy/MESRI-Finnish Society of Sciences and Letters; this study has been funded by the Agence Nationale de la Recherche (« HSF-EPISAME », SAMENTA ANR-13-SAMA-0008-01; VM).

## References

Aceros H, Der Sarkissian S, Borie M, Stevens LM, Mansour S, Noiseux N (2019) Celastrol-type HSP90 modulators allow for potent cardioprotective effects. Life Sci 227:8–19. https://doi.org/S0024-3205(19)30281-4 [pii]

Ahlskog JK, Björk JK, Elsing AN, Aspelin C, Kallio M, Roos-Mattjus P, Sistonen L (2010) Anaphase-promoting complex/cyclosome participates in the acute response to protein-damaging stress. Mol Cell Biol 30:5608–5620. https://doi.org/10.1128/MCB.01506-09 [doi]

Åkerfelt M, Morimoto RI, Sistonen L (2010) Heat shock factors: integrators of cell stress, development and lifespan. Nat Rev Mol Cell Biol 11:545–555. https://doi.org/10.1038/nrm2938 [doi]

Alastalo TP, Hellesuo M, Sandqvist A, Hietakangas V, Kallio M, Sistonen L (2003) Formation of nuclear stress granules involves HSF2 and coincides with the nucleolar localization of Hsp70. J Cell Sci 116:3557–3570. https://doi.org/10.1242/jcs.00671 [doi]

Anckar J, Hietakangas V, Denessiouk K, Thiele DJ, Johnson MS, Sistonen L (2006) Inhibition of DNA binding by differential sumoylation of heat shock factors. Mol Cell Biol 26:955–964. https://doi.org/26/3/955 [pii]

Asano J, Chiba K, Tada M, Yoshii T (1996) Cytotoxic xanthones from Garcinia hanburyi. Phytochemistry 41:815–820. https://doi.org/0031-9422(95)00682-6 [pii]

Bagatell R, Paine-Murrieta GD, Taylor CW, Pulcini EJ, Akinaga S, Benjamin IJ, Whitesell L (2000) Induction of a heat shock factor 1-dependent stress response alters the cytotoxic activity of hsp90-binding agents. Clin Cancer Res 6:3312–3318

Banik K, Harsha C, Bordoloi D, Lalduhsaki Sailo B, Sethi G, Leong HC, Arfuso F, Mishra S, Wang L, Kumar AP, Kunnumakkara AB (2018) Therapeutic potential of gambogic acid, a caged xanthone, to target cancer. Cancer Lett 416:75–86. https://doi.org/S0304-3835(17)30786-3 [pii]

Biamonti G, Vourc’h C (2010) Nuclear stress bodies. Cold Spring Harb Perspect Biol 2:a000695. https://doi.org/10.1101/cshperspect.a000695 [doi]

Björk JK, Åkerfelt M, Joutsen J, Puustinen MC, Cheng F, Sistonen L, Nees M (2016) Heat-shock factor 2 is a suppressor of prostate cancer invasion. Oncogene 35:1770–1784. https://doi.org/10.1038/onc.2015.241 [doi]

Björk JK, Sistonen L (2010) Regulation of the members of the mammalian heat shock factor family. FEBS J 277:4126–4139. https://doi.org/10.1111/j.1742-4658.2010.07828.x [doi]

Budzynski MA, Puustinen MC, Joutsen J, Sistonen L (2015) Uncoupling Stress-Inducible Phosphorylation of Heat Shock Factor 1 from Its Activation. Mol Cell Biol 35:2530–2540. https://doi.org/10.1128/MCB.00816-14 [doi]

Cascao R, Fonseca JE, Moita LF (2017) Celastrol: A Spectrum of Treatment Opportunities in Chronic Diseases. Front Med (Lausanne) 4:69. https://doi.org/10.3389/fmed.2017.00069 [doi]

Chen SR, Dai Y, Zhao J, Lin L, Wang Y, Wang Y (2018) A Mechanistic Overview of Triptolide and Celastrol, Natural Products from Tripterygium wilfordii Hook F. Front Pharmacol 9:104. https://doi.org/10.3389/fphar.2018.00104 [doi]

Chi Y, Zhan XK, Yu H, Xie GR, Wang ZZ, Xiao W, Wang YG, Xiong FX, Hu JF, Yang L, Cui CX, Wang JW (2013) An open-labeled, randomized, multicenter phase IIa study of gambogic acid injection for advanced malignant tumors. Chin Med J (Engl) 126:1642–1646

Csermely P, Schnaider T, Soti C, Prohaszka Z, Nardai G (1998) The 90-kDa molecular chaperone family: structure, function, and clinical applications. A comprehensive review. Pharmacol Ther 79:129–168. https://doi.org/S0163725898000138 [pii]

Dai C (2018) The heat-shock, or HSF1-mediated proteotoxic stress, response in cancer: from proteomic stability to oncogenesis. Philos Trans R Soc Lond B Biol Sci 373:10.1098/rstb.2016.0525. https://doi.org/20160525 [pii]

Dal Piaz F, Terracciano S, De Tommasi N, Braca A (2015) Hsp90 Activity Modulation by Plant Secondary Metabolites. Planta Med 81:1223–1239. https://doi.org/10.1055/s-0035-1546251 [doi]

Davenport J, Manjarrez JR, Peterson L, Krumm B, Blagg BS, Matts RL (2011) Gambogic acid, a natural product inhibitor of Hsp90. J Nat Prod 74:1085–1092. https://doi.org/10.1021/np200029q [doi]

Dayalan Naidu S, Dinkova-Kostova AT (2017) Regulation of the mammalian heat shock factor 1. FEBS J 284:1606–1627. https://doi.org/10.1111/febs.13999 [doi]

Duan D, Zhang B, Yao J, Liu Y, Sun J, Ge C, Peng S, Fang J (2014) Gambogic acid induces apoptosis in hepatocellular carcinoma SMMC-7721 cells by targeting cytosolic thioredoxin reductase. Free Radic Biol Med 69:15–25. https://doi.org/10.1016/j.freeradbiomed.2013.12.027 [doi]

Gomez-Pastor R, Burchfiel ET, Thiele DJ (2018) Regulation of heat shock transcription factors and their roles in physiology and disease. Nat Rev Mol Cell Biol 19:4–19. https://doi.org/10.1038/nrm.2017.73 [doi]

Han QB, Cheung S, Tai J, Qiao CF, Song JZ, Xu HX (2005) Stability and cytotoxicity of gambogic acid and its derivative, gambogoic acid. Biol Pharm Bull 28:2335–2337. https://doi.org/JST.JSTAGE/bpb/28.2335 [pii]

Hatami E, Jaggi M, Chauhan SC, Yallapu MM (2020) Gambogic acid: A shining natural compound to nanomedicine for cancer therapeutics. Biochim Biophys Acta Rev Cancer 1874:188381. https://doi.org/S0304-419X(20)30100-1 [pii]

Hentze N, Le Breton L, Wiesner J, Kempf G, Mayer MP (2016) Molecular mechanism of thermosensory function of human heat shock transcription factor Hsf1. Elife 5:10.7554/eLife.11576. https://doi.org/10.7554/eLife.11576 [doi]

Huang LE, Zhang H, Bae SW, Liu AY (1994) Thiol reducing reagents inhibit the heat shock response. Involvement of a redox mechanism in the heat shock signal transduction pathway. J Biol Chem 269:30718–30725

Huang T, Zhang H, Wang X, Xu L, Jia J, Zhu X (2019) Gambogenic acid inhibits the proliferation of smallcell lung cancer cells by arresting the cell cycle and inducing apoptosis. Oncol Rep 41:1700–1706. https://doi.org/10.3892/or.2018.6950 [doi]

Jaeger AM, Stopfer L, Lee S, Gaglia G, Sandel D, Santagata S, Lin NU, Trepel JB, White F, Jacks T, Lindquist S, Whitesell L (2019) Rebalancing Protein Homeostasis Enhances Tumor Antigen Presentation. Clin Cancer Res 25:6392–6405. https://doi.org/10.1158/1078-0432.CCR-19-0596 [doi]

Johnson JL (2012) Evolution and function of diverse Hsp90 homologs and cochaperone proteins. Biochim Biophys Acta 1823:607–613. https://doi.org/10.1016/j.bbamcr.2011.09.020 [doi]

Jolly C, Metz A, Govin J, Vigneron M, Turner BM, Khochbin S, Vourc’h C (2004) Stress-induced transcription of satellite III repeats. J Cell Biol 164:25–33. https://doi.org/10.1083/jcb.200306104 [doi]

Jolly C, Konecny L, Grady DL, Kutskova YA, Cotto JJ, Morimoto RI, Vourc’h C (2002) In vivo binding of active heat shock transcription factor 1 to human chromosome 9 heterochromatin during stress. J Cell Biol 156:775–781. https://doi.org/10.1083/jcb.200109018 [doi]

Joutsen J, Da Silva AJ, Luoto JC, Budzynski MA, Nylund AS, de Thonel A, Concordet JP, Mezger V, Saberan-Djoneidi D, Henriksson E, Sistonen L (2020) Heat Shock Factor 2 Protects against Proteotoxicity by Maintaining Cell-Cell Adhesion. Cell Rep 30:583-597.e6. https://doi.org/S2211-1247(19)31694-8 [pii]

Joutsen J, Sistonen L (2019) Tailoring of Proteostasis Networks with Heat Shock Factors. Cold Spring Harb Perspect Biol 11:10.1101/cshperspect.a034066. https://doi.org/a034066 [pii]

Kashyap D, Mondal R, Tuli HS, Kumar G, Sharma AK (2016) Molecular targets of gambogic acid in cancer: recent trends and advancements. Tumour Biol 37:12915–12925. https://doi.org/10.1007/s13277-016-5194-8 [doi]

Kijima T, Prince TL, Tigue ML, Yim KH, Schwartz H, Beebe K, Lee S, Budzynski MA, Williams H, Trepel JB, Sistonen L, Calderwood S, Neckers L (2018) HSP90 inhibitors disrupt a transient HSP90-HSF1 interaction and identify a noncanonical model of HSP90-mediated HSF1 regulation. Sci Rep 8:6976–w. https://doi.org/10.1038/s41598-018-25404-w [doi]

Kmiecik SW, Le Breton L, Mayer MP (2020) Feedback regulation of heat shock factor 1 (Hsf1) activity by Hsp70-mediated trimer unzipping and dissociation from DNA. EMBO J 39:e104096. https://doi.org/10.15252/embj.2019104096 [doi]

Lee JH, Koo TH, Yoon H, Jung HS, Jin HZ, Lee K, Hong YS, Lee JJ (2006) Inhibition of NF-kappa B activation through targeting I kappa B kinase by celastrol, a quinone methide triterpenoid. Biochem Pharmacol 72:1311–1321. https://doi.org/S0006-2952(06)00504-1 [pii]

Lee JY, Lee BH, Kim ND, Lee JY (2015) Celastrol blocks binding of lipopolysaccharides to a Toll-like receptor4/myeloid differentiation factor2 complex in a thiol-dependent manner. J Ethnopharmacol 172:254–260. https://doi.org/10.1016/j.jep.2015.06.028 [doi]

Liu P, Wu X, Dai L, Ge Z, Gao C, Zhang H, Wang F, Zhang X, Chen B (2017) Gambogenic Acid Exerts Antitumor Activity in Hypoxic Multiple Myeloma Cells by Regulation of miR-21. J Cancer 8:3278–3286. https://doi.org/10.7150/jca.19290 [doi]

Mahat DB, Salamanca HH, Duarte FM, Danko CG, Lis JT (2016) Mammalian Heat Shock Response and Mechanisms Underlying Its Genome-wide Transcriptional Regulation. Mol Cell 62:63–78. https://doi.org/10.1016/j.molcel.2016.02.025 [doi]

Masser AE, Ciccarelli M, Andreasson C (2020) Hsf1 on a leash - Controlling the heat shock response by chaperone titration. Exp Cell Res:112246. https://doi.org/S0014-4827(20)30495-X [pii]

Mathew A, Mathur SK, Morimoto RI (1998) Heat shock response and protein degradation: regulation of HSF2 by the ubiquitin-proteasome pathway. Mol Cell Biol 18:5091–5098. https://doi.org/10.1128/mcb.18.9.5091 [doi]

Neckers L, Workman P (2012) Hsp90 molecular chaperone inhibitors: are we there yetã. Clin Cancer Res 18:64–76. https://doi.org/10.1158/1078-0432.CCR-11-1000 [doi]

Östling P, Bjork JK, Roos-Mattjus P, Mezger V, Sistonen L (2007) Heat shock factor 2 (HSF2) contributes to inducible expression of hsp genes through interplay with HSF1. J Biol Chem 282:7077–7086. https://doi.org/M607556200 [pii]

Palempalli UD, Gandhi U, Kalantari P, Vunta H, Arner RJ, Narayan V, Ravindran A, Prabhu KS (2009) Gambogic acid covalently modifies IkappaB kinase-beta subunit to mediate suppression of lipopolysaccharide-induced activation of NF-kappaB in macrophages. Biochem J 419:401–409. https://doi.org/10.1042/BJ20081482 [doi]

Park SM, Kim SA, Ahn SG (2015) HSF2 autoregulates its own transcription. Int J Mol Med 36:1173–1179. https://doi.org/10.3892/ijmm.2015.2309 [doi]

Pearl LH (2016) Review: The HSP90 molecular chaperone-an enigmatic ATPase. Biopolymers 105:594–607. https://doi.org/10.1002/bip.22835 [doi]

Peng B, Xu L, Cao F, Wei T, Yang C, Uzan G, Zhang D (2010) HSP90 inhibitor, celastrol, arrests human monocytic leukemia cell U937 at G0/G1 in thiol-containing agents reversible way. Mol Cancer 9:79–79. https://doi.org/10.1186/1476-4598-9-79 [doi]

Puustinen MC, Sistonen L (2020) Molecular Mechanisms of Heat Shock Factors in Cancer. Cells 9:10.3390/cells9051202. https://doi.org/E1202 [pii]

Richter K, Haslbeck M, Buchner J (2010) The heat shock response: life on the verge of death. Mol Cell 40:253–266. https://doi.org/10.1016/j.molcel.2010.10.006 [doi]

Salminen A, Lehtonen M, Paimela T, Kaarniranta K (2010) Celastrol: Molecular targets of Thunder God Vine. Biochem Biophys Res Commun 394:439–442. https://doi.org/10.1016/j.bbrc.2010.03.050 [doi]

Sandqvist A, Björk JK, Åkerfelt M, Chitikova Z, Grichine A, Vourc’h C, Jolly C, Salminen TA, Nymalm Y, Sistonen L (2009) Heterotrimerization of heat-shock factors 1 and 2 provides a transcriptional switch in response to distinct stimuli. Mol Biol Cell 20:1340–1347. https://doi.org/10.1091/mbc.E08-08-0864 [doi]

Santagata S, Xu YM, Wijeratne EM, Kontnik R, Rooney C, Perley CC, Kwon H, Clardy J, Kesari S, Whitesell L, Lindquist S, Gunatilaka AA (2012) Using the heat-shock response to discover anticancer compounds that target protein homeostasis. ACS Chem Biol 7:340–349. https://doi.org/10.1021/cb200353m [doi]

Sarge KD, Murphy SP, Morimoto RI (1993) Activation of heat shock gene transcription by heat shock factor 1 involves oligomerization, acquisition of DNA-binding activity, and nuclear localization and can occur in the absence of stress. Mol Cell Biol 13:1392–1407. https://doi.org/10.1128/mcb.13.3.1392 [doi]

Seo MJ, Lee DM, Kim IY, Lee D, Choi MK, Lee JY, Park SS, Jeong SY, Choi EK, Choi KS (2019) Gambogic acid triggers vacuolization-associated cell death in cancer cells via disruption of thiol proteostasis. Cell Death Dis 10:187–4. https://doi.org/10.1038/s41419-019-1360-4 [doi]

Shen D, Wang Y, Niu H, Liu C (2020) Gambogenic acid exerts anticancer effects in cisplatinresistant nonsmall cell lung cancer cells. Mol Med Rep 21:1267–1275. https://doi.org/10.3892/mmr.2020.10909 [doi]

Shrestha L, Bolaender A, Patel HJ, Taldone T (2016) Heat Shock Protein (HSP) Drug Discovery and Development: Targeting Heat Shock Proteins in Disease. Curr Top Med Chem 16:2753–2764. https://doi.org/CTMC-EPUB-74968 [pii]

Sreedhar AS, Kalmar E, Csermely P, Shen YF (2004) Hsp90 isoforms: functions, expression and clinical importance. FEBS Lett 562:11–15. https://doi.org/10.1016/s0014-5793(04)00229-7 [doi]

Tang Q, Lu M, Zhou H, Chen D, Liu L (2017) Gambogic acid inhibits the growth of ovarian cancer tumors by regulating p65 activity. Oncol Lett 13:384–388. https://doi.org/10.3892/ol.2016.5433 [doi]

Trott A, West JD, Klaic L, Westerheide SD, Silverman RB, Morimoto RI, Morano KA (2008) Activation of heat shock and antioxidant responses by the natural product celastrol: transcriptional signatures of a thiol-targeted molecule. Mol Biol Cell 19:1104–1112. https://doi.org/10.1091/mbc.E07-10-1004 [doi]

Vihervaara A, Mahat DB, Guertin MJ, Chu T, Danko CG, Lis JT, Sistonen L (2017) Transcriptional response to stress is pre-wired by promoter and enhancer architecture. Nat Commun 8:255–0. https://doi.org/10.1038/s41467-017-00151-0 [doi]

Vihervaara A, Sergelius C, Vasara J, Blom MA, Elsing AN, Roos-Mattjus P, Sistonen L (2013) Transcriptional response to stress in the dynamic chromatin environment of cycling and mitotic cells. Proc Natl Acad Sci U S A 110:3388. https://doi.org/10.1073/pnas.1305275110 [doi]

Vuori KA, Ahlskog JK, Sistonen L, Nikinmaa M (2009) TransLISA, a novel quantitative, nonradioactive assay for transcription factor DNA-binding analyses. FEBS J 276:7366–7374. https://doi.org/10.1111/j.1742-4658.2009.07446.x [doi]

Wang Y, McAlpine SR (2015) Heat-shock protein 90 inhibitors: will they ever succeed as chemotherapeuticsã. Future Med Chem 7:87–90. https://doi.org/10.4155/fmc.14.154 [doi]

Wang Y, Gibney PA, West JD, Morano KA (2012) The yeast Hsp70 Ssa1 is a sensor for activation of the heat shock response by thiol-reactive compounds. Mol Biol Cell 23:3290–3298. https://doi.org/10.1091/mbc.E12-06-0447 [doi]

Weerapana E, Simon GM, Cravatt BF (2008) Disparate proteome reactivity profiles of carbon electrophiles. Nat Chem Biol 4:405–407. https://doi.org/10.1038/nchembio.91 [doi]

Westerheide SD, Anckar J, Stevens SM, Sistonen L, Morimoto RI (2009) Stress-inducible regulation of heat shock factor 1 by the deacetylase SIRT1. Science 323:1063–1066. https://doi.org/10.1126/science.1165946 [doi]

Westerheide SD, Bosman JD, Mbadugha BN, Kawahara TL, Matsumoto G, Kim S, Gu W, Devlin JP, Silverman RB, Morimoto RI (2004) Celastrols as inducers of the heat shock response and cytoprotection. J Biol Chem 279:56053–56060. https://doi.org/M409267200 [pii]

Wu ZQ, Guo QL, You QD, Zhao L, Gu HY (2004) Gambogic acid inhibits proliferation of human lung carcinoma SPC-A1 cells in vivo and in vitro and represses telomerase activity and telomerase reverse transcriptase mRNA expression in the cells. Biol Pharm Bull 27:1769–1774. https://doi.org/JST.JSTAGE/bpb/27.1769 [pii]

Xia G, Wang H, Song Z, Meng Q, Huang X, Huang X (2017) Gambogic acid sensitizes gemcitabine efficacy in pancreatic cancer by reducing the expression of ribonucleotide reductase subunit-M2 (RRM2). J Exp Clin Cancer Res 36:107–0. https://doi.org/10.1186/s13046-017-0579-0 [doi]

Yang J, Li C, Ding L, Guo Q, You Q, Jin S (2012) Gambogic acid deactivates cytosolic and mitochondrial thioredoxins by covalent binding to the functional domain. J Nat Prod 75:1108–1116. https://doi.org/10.1021/np300118c [doi]

Yim KH, Prince TL, Qu S, Bai F, Jennings PA, Onuchic JN, Theodorakis EA, Neckers L (2016) Gambogic acid identifies an isoform-specific druggable pocket in the middle domain of Hsp90beta. Proc Natl Acad Sci U S A 113:4801. https://doi.org/10.1073/pnas.1606655113 [doi]

Yuno A, Lee MJ, Lee S, Tomita Y, Rekhtman D, Moore B, Trepel JB (2018) Clinical Evaluation and Biomarker Profiling of Hsp90 Inhibitors. Methods Mol Biol 1709:423–441. https://doi.org/10.1007/978-1-4939-7477-1_29 [doi]

Zhou J, Luo YH, Wang JR, Lu BB, Wang KM, Tian Y (2013) Gambogenic acid induction of apoptosis in a breast cancer cell line. Asian Pac J Cancer Prev 14:7601–7605. https://doi.org/10.7314/apjcp.2013.14.12.7601 [doi]

Zou J, Guo Y, Guettouche T, Smith DF, Voellmy R (1998) Repression of heat shock transcription factor HSF1 activation by HSP90 (HSP90 complex) that forms a stress-sensitive complex with HSF1. Cell 94:471–480. https://doi.org/S0092-8674(00)81588-3 [pii]

